# CORSICA: Reproducible Suppression of Cochlear Implant Artifacts in EEG evoked by Continuous Speech

**DOI:** 10.64898/2026.07.27.740877

**Authors:** Constantin Jehn, Clara Stiller, Niki Katerina Vavatzanidis, Tobias Reichenbach

**Affiliations:** Department Artificial Intelligence in Biomedical Engineering (AIBE), Friedrich-Alexander-Universität Erlangen-Nürnberg (FAU), Nürnberger Strasse 74, 91052 Erlangen, Germany; Ear Research Center Dresden (ERCD), University Hospital Carl Gustav Carus, Technische Universität Dresden, Dresden, Germany

## Abstract

**Objective:** Electroencephalography (EEG) is a key tool for studying auditory processing in cochlear implant (CI) users. In particular, EEG recordings obtained during continuous speech are becoming increasingly important for assessing speech and language processing in CI users, and may be utilized for neurofeedback. However, CIs also induce strong stimulation artifacts that are time-locked to the stimulus and mask the neural responses that have smaller magnitudes. Existing artifact reduction methods are typically based on event-related potentials (ERPs) or require manual component selection, making them unsuitable for naturalistic listening conditions or large datasets.

**Approach:** We develop CORSICA (CORrelation-baSed ICA artifact rejection), a reproducible, parameter-efficient method for CI artifact reduction in EEG responses to continuous speech. CORSICA operates on independent components (ICs) obtained through Infomax ICA and requires no manual component labelling, with performance governed by a single tunable threshold. It exploits the observation that CI artifacts temporally follow the audio signal without delay, whereas neural responses have an inherent lag due to auditory pathway latencies. For each IC, CORSICA computes the cross-correlation with the speech stimulus. Artifacts are identified by a high signal-to-noise ratio (SNR) of the correlation peak near zero lag, and the component is rejected if this SNR exceeds a threshold. To benchmark CORSICA, we evaluate two alternatives: a TRF-based SNR method, in which temporal response functions are fitted to each IC and artifact-driven peaks near zero lag are used for rejection, and a variant replacing ICA with second-order blind identification (SOBI) as the source separation step.

**Main results:** CORSICA effectively suppressed CI artifacts while preserving neural activity, enabling recovery of physiologically plausible TRFs with only 2% of ICs rejected. Both benchmark methods confirmed the validity of the SNR-based rejection framework, but CORSICA outperformed the TRF-based alternative in artifact suppression quality. Replacing ICA with SOBI as the source separation step required more ICs to be rejected, further supporting ICA as the preferred backbone for CORSICA.

**Significance:** CORSICA provides a fully objective, label-free approach to identifying CI artifacts in speech-evoked EEG data, with no manual intervention required. By centering artifact rejection on a single interpretable threshold, it offers a reproducible preprocessing standard for future EEG studies on speech processing in CI users.

**Conclusion:** Our findings demonstrate that objective CI artifact suppression in speech-evoked EEG data is feasible on the basis of the IC’s temporal response patterns.

## 1 Introduction

Cochlear implants (CIs) have significantly advanced the treatment of severe to profound hearing loss by directly stimulating the auditory nerve, allowing individuals to regain access to sound and, most importantly, to speech. In a CI system, acoustic signals are captured by an external microphone, converted into electrical stimuli, and transmitted to the implanted receiver. The signals are then decoded into pulse trains that stimulate the auditory nerve through an electrode inserted into the cochlea [1, 2]. Various signal coding strategies are employed for this transformation, typically optimized for speech signals to enhance speech perception and intelligibility [3].

Reliable electroencephalography (EEG) measurements of brain activity in CI users are essential for both clinical and research applications. EEG-based assessments can provide objective evaluations of implant function shortly after surgery [4], supporting outcome evaluation and device fitting [4], particularly in young children who cannot report their hearing perception [5]. Furthermore, EEG can allow to study speech processing in CI users, particularly regarding speech perception in complex listening environments requiring selective attention, such as multi-talker scenarios [6, 7, 8, 9]. Finally, EEG recordings could be used for neurofeedback to the CI. As an example, when faced with multiple competing speakers, EEG recordings can allow to decode the user’s focus of attention, and feed this information back to the CI [10, 11]. The device may then enhance the target speech stream to facilitate its understanding for the user, potentially improving speech comprehension and listening comfort [12].

However, EEG recordings in CI users are typically contaminated by electrical artifacts induced by the implant’s stimulation, presenting a significant technical challenge. These artifacts often exhibit amplitudes much greater than the underlying neural signals and are time-locked to the stimulus [13], complicating the separation of neural responses from artifacts. Additionally, the characteristics of CI artifacts and how these manifest in EEG recordings can highly vary across individuals, making artifact identification challenging [14].

Several methods have been developed to reduce CI-induced artifacts in EEG recordings. A widely adopted approach is blind source separation using independent component analysis (ICA), which decomposes EEG signals into statistically independent components (ICs), each representing a distinct source that contributes to the recorded signal [15]. The ICs can then be manually inspected for the presence of artifacts, guided by known artifact characteristics [7, 13, 14]. The characteristics are (1) CI artifacts align with stimulus onset and offset (including filter ringing), (2) they persist throughout stimulus presentation, and (3) they exhibit scalp topographies centered near the implant site [13].

To reduce manual inspection requirements, Viola et al. proposed a semi-automatic method for the identification of artifact components [14]. Their approach selects a template IC that best represents the artifact by correlating it with all other ICs from the same subject, identifying highly correlated components for removal. Template selection is based on two criteria. Firstly, spatial characteristics that are evaluated through dipole fitting: CI artifacts show topographies that are poorly captured by dipole fits. Secondly, temporal features: CI artifacts exhibit sharp onsets and offsets that are aligned with onsets and offsets in the stimulus. This ICA-based method has shown effectiveness for analyzing event-related potentials (ERPs) with short-duration stimuli, such as individual syllables like /ba/ lasting around 100 ms. Moreover, Deprez et al. proposed to classify independent components based on their spectral power to measure clean evoked auditory steady-state responses (i.e., responses to periodic stimulation from the IC) that also work with epoched data [16]. This approach assumes that only CI artifacts have high-frequency components above 200 Hz and can thus be separated from brain responses. Recent years, however, have seen growing research interest in studying neural responses to continuous speech in CI users [5, 7]. This approach permits the investigation of neural processing of speech in more natural listening conditions, including situations such as continuous speech in background noise that CI users find challenging. However, longer and continuous stimuli introduce additional challenges for artifact suppression, as neural responses and artifacts overlap persistently throughout the stimulus duration. Moreover, complex stimuli like speech generate artifacts of lower magnitude compared to simpler stimuli, such as pure tones, due to the wide-ranging and more variable amplitude of the speech signals [1]. The way these amplitude modulations of the artifact manifest in actual EEG recordings depends on user-specific CI configurations and settings, as well as implant location, and therefore varies across individuals [1]. Last but not least, the approach by Viola requires averaging of the EEG signal over many repeated presentations of the same sound, which is not available for EEG responses to longer continuous speech [14, 17].

To address artifact removal for EEG responses to continuous speech, alternative methods have been proposed. One approach involves inserting periodic gaps in the stimulation to allow artifact decay, enabling measurement of uncontaminated neural responses during these gaps [10]. This method uses 4 ms stimulation gaps every 40 ms, successfully reducing artifacts without compromising speech intelligibility. However, this approach requires stimulus modification, making it impractical in natural listening conditions.

Another approach suggested by Paul et al. uses blind source separation for continuous speech and manual identification of artifact components [7]. This method employs second-order blind identification (SOBI), which separates temporally correlated sources, in contrast to ICA’s assumption of statistical independence between sources. This method accounts for the temporal correlation of CI artifacts with the stimulus, allowing separation of CI artifact components from neural signals. Components’ topographies matching the spatial location of CI implants are further analyzed by visually inspecting their time course with the envelope of the audio stimulus. Components showing high temporal similarity to the stimulus are removed. Furthermore, Althoff and Nogueira proposed learning the weights of the backward model, i.e. a model that reconstructs the presented stimulus from the recorded neural data, directly on the time series of independent components derived via SOBI [18]. This allowed the backward model to down-weight artifactual components that were irrelevant to the reconstruction task. The hope was that this leads to a data-driven suppression of artifacts. The method was validated on a selective listening task in which participants focused on one of two concurrently playing instruments. This approach reduced the high reconstruction scores that were caused by the CI artifacts at early latencies (*<* 100 ms) to the level of typical hearing individuals, proving its usefulness in obtaining interpretable results from the backward models. However, the method did not significantly improve decoding accuracy. Its efficacy was not tested on forward models that are more susceptible to CI artifacts.

Despite these advances and the growing interest in studying EEG responses to continuous speech in CI users, a systematic evaluation of CI artifact reduction methods is lacking. We have recently proposed classifying ICs as CI artifacts based on their temporal correlation to the stimulus [11]: since CI-induced artifacts occur at the short delay of CI processing, components with a high correlation SNR near 0 ms are flagged for rejection. The present study formalizes and extends this approach into CORSICA (CORrelation-baSed ICA artifact rejection). Specifically, we broaden the artifact search window to improve detection sensitivity and provide the first thorough benchmarking of the method. As reference, we evaluate a TRF-based SNR method as an alternative rejection criterion, as well as SOBI as an alternative to ICA for source separation. The latter is motivated by its growing popularity in CI artifact detection. All approaches are evaluated on their ability to recover clean, physiologically plausible TRFs, which serve as a sensitive readout of residual CI contamination.

## 2 Methods

### 2.1 Data set

We used a data set that we recorded for a previous study [9], and that is publicly available [19]. Below, we summarize the main aspects of this experiment.

#### 2.1.1 Participants

Twenty-four bilaterally implanted CI users were assessed. All participants were native German speakers without cognitive impairment. CI experience ranged from 2 to 28 years (median: 10 years). Inclusion required a minimum score of 60% on the Freiburg monosyllabic word recognition test in quiet at 65 dB for the better-performing ear [20], ensuring sufficient speech perception for the selective attention paradigm. Speech understanding in noise was further assessed using the HSM sentence test 65 dB signal, 60 dB noise, free field) [21]. Most CI users had post-lingual hearing loss (n=20), while four were classified as pre- or peri-lingual. Detailed information about the participants and their implants, including processor and stimulation strategy was previously reported in [9], where it can be found in the Appendix.

The experiment was approved by the science ethics committee of Technische Universität Dresden (protocol SR+BO-EK-47022024) on May 21, 2024. All procedures were conducted in accordance with the Declaration of Helsinki.

#### 2.1.2 Experiment and stimuli

The experiment consisted of an auditory attention task. Excerpts from two German audiobooks were used as auditory stimuli: *Elbenwald - Blatt von Tüftler* narrated by a male voice, and *Eine Frau erlebt die Polarnacht*, narrated by a female voice. The audiobook excerpts were segmented into trials with an average duration of two minutes and presented at an individually selected, comfortable loudness level using each participant’s standard CI settings.

The audiobooks were presented over two loudspeakers separated by an angle of 60°, located in front of the participants, in a free-field environment. The experiment comprised two scenarios. The first eight trials were in a single-speaker condition, in which only one audiobook was played through one of the two speakers. The following twelve trials were in a competing speaker condition, during which both audiobooks were played simultaneously, each story from a separate loudspeaker. Participants were instructed to focus on one speaker at a time, with the focus of attention changing from trial to trial. After each trial, participants answered comprehension questions to validate their attentiveness to the target speaker.

#### 2.1.3 EEG recordings

EEG data were recorded using a 32-channel actiCHamp system (BrainProducts GmbH, Germany) at a sampling frequency of 1 kHz with electrode Cz as online reference. The signal was online low-pass filtered with a cut-off frequency of 280 Hz. Electrode impedances were checked before the recording and during the break between the single and competing speaker conditions. They were kept below 20 *k*Ω. Up to two electrodes per hemisphere were removed due to close positioning relative to the CI coil, making reliable recording at these sites infeasible. On average, 1.7 electrodes per participant were removed across both hemispheres. Audio signals were recorded over two auxiliary channels, using two StimTrak adapters (BrainProducts GmbH, Germany), to align EEG data and audio stimuli.

### 2.2 Preprocessing of EEG Data

EEG data preprocessing was performed in MATLAB using the EEGLAB toolbox. The main steps are illustrated in Fig. 1A. For synchronization, the presented audio was recorded as auxiliary channel via the EEG amplifier using StimTrak adapters (BrainProducts GmbH, Germany). Alignment between the EEG and each audio stimulus was then achieved offline by cross-correlating the recorded audio with delayed versions of the clean stimulus and selecting the delay that maximized Pearson’s correlation coefficient. EEG data were then segmented into single trials. Channels that had been removed before the measurement due to their proximity to the implant were interpolated to maintain dimension consistency across datasets. Furthermore, an average reference was applied. Baseline drift was removed by applying a high-pass filter using a sinc zero-phase FIR filter, introducing no temporal delay, with a Hamming window at a cutoff frequency of 1 Hz.

**Figure 1:**
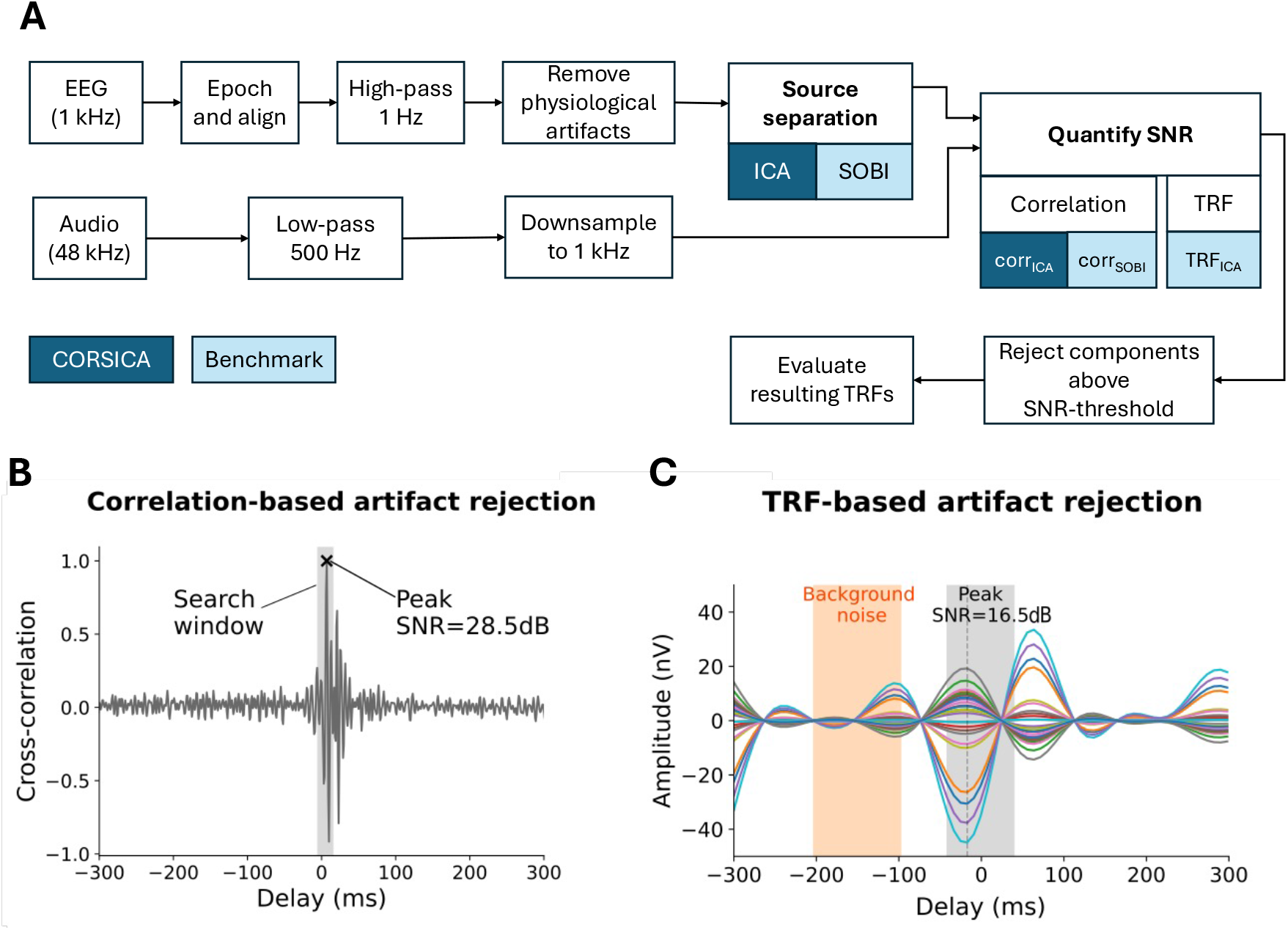
Overview of the proposed CORSICA method. **A**, EEG and audio processing pipeline. Following standard pre-processing (high-pass filtering and physiological artifact removal), source separation is performed using either ICA or SOBI (the latter as a benchmark alternative). The audio signal is low-pass filtered at 500 Hz and downsampled to 1 kHz. The SNR of each source relative to the audio is then quantified via cross-correlation or TRFs. Methods for CI artifact detection are named by combining the source separation technique with the SNR quantification method: corr_ICA_ (correlation-based SNR quantification after ICA, referred to as CORSICA throughout the text), corr_SOBI_ (correlation-based SNR quantification after SOBI), and TRF_ICA_ (TRF-based SNR quantification after ICA). In all cases, components above an SNR threshold are excluded. **B**, For the correlation-based method the cross-correlation between the speech stimulus and an independent component of the EEG is computed. The exemplary cross-correlation shows a clear peak in a search window around 0 ms delay (delays of *−*5 – 12 ms, gray shading). The SNR of this peak is computed from the peak amplitude and the average correlation outside the search window. The highlighted peak in the search window has an SNR of 28.5 dB. **C**, For the TRF-based method, TRFs are computed for each IC of the EEG recordings to the sum of the attended and ignored speech envelope. The peak within the light gray time window (0 ms *±* 40 ms) is identified. The exemplary TRFs show a distinct peak in this search area. The SNR value is calculated from the ratio of the peak value and a noise estimate, defined as the average TRF amplitude at negative time lag, between *−*200 and *−*100 ms. The exemplary TRF reaches an SNR of 16.5 dB.

To remove physiological artifacts, InfoMax ICA was applied at the trial level to separate up to 30 components, or fewer if the rank of the data was lower, and automatic classification using ICLabel [22] was applied. Components labeled as muscle, ocular, or cardiac artifact with a probability of greater than 60% were excluded, which led to an average exclusion of 27.1% of components (range per participant: 16.1% – 38.9%). After removing these physiological artifacts, we then applied the proposed strategies for CI-artifact removal.

For the subsequent TRF-analysis, EEG data were low-pass filtered with an upper passband edge of 8 Hz (one-pass, zero-phase, non-causal type 1 FIR filter, −6 dB cutoff frequency: 9.0 Hz, order 1,650, Hamming window with 0.0194 passband ripple and 53 dB stopband attenuation) and then downsampled to 125 Hz.

### 2.3 Temporal Response Functions

We assessed the neural tracking of the speech envelope through computing Temporal Response Functions (TRFs) [23, 24]. In particular, a linear forward model was employed to reconstruct the EEG response from the speech envelope of the attended and of the distractor speaker separately. The model coefficients at the different delays constitute the TRF. Their morphology yields insight into the spatio-temporal neural response to the stimulus [25]. Because the neural response to the speech envelope is generally well characterized in typical hearing listeners, CI artifacts in the TRFs can often be identified. Although their precise morphology may vary across CI users, salient features such as persistent and large early peaks are typically indicative of residual CI artifacts, and TRFs can thus be employed to evaluate the effectiveness of CI-artifact suppression.

To extract the speech envelope of the audio stimuli, we applied the Hilbert transform to the audiobooks, followed by taking the absolute value of the resulting signal. Afterwards, the envelopes were bandpass-filtered between 1 and 8 Hz using a linear-phase FIR filter with a Hamming window. The data was then downsampled to 125 Hz and normalized using the z-score transformation.

TRFs were computed using the sPyEEG package in Python, which implements ridge regression for regularized linear estimation [26]. To avoid the confounding influence of different regularization coefficients on the TRF shape, the regularization parameter was set to 1 for all TRF calculations. The analysis window of temporal delays ranged from −300 ms to 500 ms. Negative time lags were included to verify that no neural responses emerged that preceded the stimulation. The TRFs were computed on single-trial data of the competing speaker paradigm. The resulting model coefficients were averaged across trials and participants to obtain grand-average TRFs.

Beyond their role in characterizing auditory processing, TRFs computed on individual independent components also serve as a criterion for artifact identification, as described in Subsection 2.4, and as the primary evaluation subject for comparing artifact reduction methods, as described in Subsections 2.5 and 2.6.

### 2.4 CI artifact reduction

To identify and remove independent components contaminated by CI artifacts, we employed CORSICA, which quantifies the relationship between the audio stimulus and each IC via cross-correlation. We contrasted the results with those obtained by a TRF-based approach that served as a benchmark.

Both methods exploit the observation that CI-induced artifacts mirror the audio stimulus with a minimal temporal delay of up to 10 ms for bilaterally implanted CIs [27], and classify an IC as artifactual if it exhibits this characteristic near-zero-latency relationship with the stimulus. In contrast, neural responses to the speech envelope are delayed by synaptic and neural conduction times along the auditory pathway, with the first EEG response to the speech envelope typically measured between 30–80 ms after stimulus onset [28].

Prior to artifact detection, the audio signal was low-pass filtered below 500 Hz and resampled to 1 kHz to match the EEG signal, as depicted in Fig. 1A.

Crucially, source separation was performed at the trial level rather than across the entire recording, providing a fine-grained resolution that allowed artifact characteristics to be captured within individual trials. Both methods were implemented in Python.

#### 2.4.1 CORSICA

CORSICA builds on the correlation-based artifact detection method proposed in [11], which is illustrated in Fig. 1B. Here, we refine this method and assess its performance systematically through comparison to two other benchmark methods. In particular, as a methodological refinement, we broaden the search window around zero delay, which improves detection sensitivity for CI artifact peaks.

A full cross-correlation (as implemented in `scipy.signal.correlate` with `mode=’full’`) between the audio stimulus, consisting of both the attended and distractor audiobook, and the time series of each independent component, separated using InfoMax ICA, was computed, yielding correlation values at all possible delays. Neural responses should not show a sizable correlation, since they occur at much lower frequencies, and if so, at positive delays due to the neural processing. In contrast, a noticeable correlation at a delay of about zero indicates a CI artifact.

To identify a potential peak in the correlation function around zero, we employed a search window between *−*5 ms and 12 ms to account for the processing delay of the CI. To quantify the artifact strength, the signal-to-noise ratio (SNR) of the largest correlation peak in the search window was calculated. The background noise was thereby defined as the root-mean-square of the cross-correlation signal outside the search window; that is, all delays availabl from the full cross-correlation that fall outside the *−*5 ms to 12 ms range. The SNR was then converted to decibels.

Components exceeding a certain SNR threshold were considered to contain mainly CI artifacts and were excluded from further analysis. The SNR threshold served as a tunable parameter in the algorithm that could be adjusted to yield optimal results. Its choice represents a trade-off between maximal artifact suppression while minimizing the loss of data. The EEG data were then reconstructed using the remaining components, yielding a dataset with reduced artifact interference.

#### 2.4.2 SOBI blind source separation

As an alternative to InfoMax ICA, second-order blind identification (SOBI) [29] has recently been employed to separate sources for subsequent CI artifact identification [7, 18]. SOBI performs joint diagonalization of time-delayed covariance matrices, which implicitly minimizes the cross-correlation between the identified components for a specified number of delays, and has been applied broadly to remove various types of artifacts in EEG, including ocular and muscle artifacts [30, 31].

To assess its effectiveness for CI artifact detection, SOBI was used to separate up to 30 sources per trial, or fewer if the data rank was lower. As longer sets of delays lead to more robust source separation for EEG [32], the maximal lag was set to 250 samples, corresponding to delays between 0 and 250 ms for the calculation of the auto-covariance matrices. After that, we applied correlation-based artifact detection method as shown in Fig. 1B. We refer to this benchmark as corr_SOBI_. To facilitate a fair comparison between the ICA-based and the SOBI-based source-separation method, we compared data sets with an equivalent number of retained IC components.

#### 2.4.3 TRF-based artifact detection

The TRF-based artifact detection method, illustrated in Fig. 1C, identifies CI artifacts by fitting TRFs from each independent component to the summed speech envelope of the attended and ignored audiobook. As with the correlation-based method, a large TRF peak near 0 ms latency is indicative of a CI artifact rather than genuine neural activity.

TRFs were computed for each independent component using the same parameters as described in 2.3. To account for temporal smearing of responses due to low-pass filtering of the envelopes and the EEG, we searched for the maximum TRF value within a broader search window of delays than for the correlation-based method, namely within ±40 ms of delay around 0 ms. We then computed the SNR for each TRF from the ratio of the average peak amplitude within this window to the background noise. The latter was quantified as the mean-square amplitude of the TRFs in a window of delays between −200 ms and −100 ms (Fig. 1B). This pre-stimulus window was chosen to ensure separation both from neural responses and from regularization effects near the smallest considered time lag of −300 ms, while also avoiding overlap with potential CI artifact peaks near 0 ms.

Components were rejected if their calculated SNR exceeded a certain threshold. As for CORSICA, the SNR threshold served as a tunable parameter in the algorithm. In the following, we refer to this TRF-based method when employing independent EEG components computed through ICA as the TRF_ICA_ method. Because we found that the TRF_ICA_-based method was inferior to CORSICA (see Results below), we did not evaluate the TRF-based method on EEG data processed through SOBI (see 2.4.2) instead of ICA.

### 2.5 Component characterization

To provide a broader characterization of CI artifact components, which has not been systematically examined before, we assessed four characteristics that give an indication of whether a component is likely a CI artifact or of neural origin.

#### Topography

Each independent component is a linear superposition of the different EEG channels. The topography of a component indicates the weights with which each channel contributes to the component. For CI artifacts, we expect topographies to be heavily lateralized and concentrated near one of the implants’ positions. In bilateral CI users, non-lateralized dipoles are in principle conceivable; however, empirical observation consistently yields strongly lateralized artifact components, likely reflecting the dominance of one implant’s electrical interference in a given component.

#### Cross-correlation with the audio

We further computed the cross-correlation function between the time-series of the ICs and the sum of the waveforms of the attended and ignored audio book. Both audio streams were included to avoid a bias in the artifact rejection stage towards either of the two speakers. As CI artifacts are highly correlated and time-locked to the stimulus, we expected pronounced peaks around 0 ms latency.

#### TRFs

Moreover, we computed each IC’s TRF to the sum of the speech envelopes of the attended and ignored audiobook, as described above. CI artifacts are indeed not only highly correlated to the waveform of the audio stimulus, but also to its envelope. Thus, we expected dominant peaks around 0 ms latency, driven by channels near the implant for artifactual components.

#### Power spectrum

We also computed how the power of representative ICs was distributed across different frequency bands using the power spectrum. For neural components, we expected peaks in the delta (1–4 Hz) and alpha (8–13 Hz) bands, and a sharp cut-off for higher frequencies. For CI-artifacts, we expected additional contributions at higher frequencies, as typical CIs stimulate up to 8 kHz [33].

### 2.6 Metrics for comparing the artifact rejection methods

To quantitatively compare the performance of the different artifact detection methods and the effect of varying the SNR thresholds, we assessed the resulting TRFs after artifact removal using metrics proposed by Kulasingham et al. [28]. Since completely artifact-free data is not available for EEG acquired from CI-users, a reference TRF was manually selected out of all the resulting TRFs (Fig. 3). Note that, this manual selection is only required for the evaluation of the effectiveness of the different methods. The artifact reduction itself remains automated with the selection of only a single hyperparameter.

This reference TRF was chosen based on (1) visually appearing free from CI artifacts, and (2) containing as much of the original EEG data as possible, and (3) corresponding to the attended speech stimulus, as the attended audio causes larger neural responses than the ignored speech signal. A TRF was considered artifact-free by visual inspection when no single channel dominated the response and the scalp topography showed a physiologically plausible pattern, such as a bipolar distribution consistent with auditory processing. The so-identified reference TRF thus represented our optimal compromise between artifact suppression and minimal data loss.

#### Pearson correlation

As the first metric, we quantified the similarity between each TRF and the reference TRF by calculating the Pearson correlation coefficient between the two TRFs for each channel, and then averaging across all channels.

#### Latency error

Peak latency accuracy was assessed by calculating the absolute timing error of the three canonical TRF components. Following established window ranges of 30–80 ms, 90–170 ms, and 190–250 ms [28, 34, 35], we identified peaks by locating the maximum root-mean-square (RMS) amplitude within each interval. The latency error followed as the mean absolute difference between the reference and reconstructed peak latencies.

#### Amplitude error

To assess signal fidelity, we calculated the mean amplitude error across the three identified peak latency windows. For each window, we measured the absolute channel-wise difference between the reconstructed and reference TRF peaks. This metric quantifies how effectively each method suppresses artifacts while preserving the original neural signal.

### 2.7 Individual and statistical analysis

We further analyzed subject-level TRFs to characterize inter-individual variability. Subject-level TRFs are inherently noisy due to the low SNR of EEG and averaging across channels and/or subjects is typically appropriate [36]. Nevertheless, variability across participants is an important aspect of both CORSICA’s performance and TRF morphology in CI users more generally.

For each subject, Pearson correlation, latency error, and amplitude error were computed against the same reference TRF used in the group-level analysis, yielding a distribution of these metrics across participants and SNR thresholds. To illustrate the range of outcomes, we additionally selected representative good, medium, and bad examples based on Pearson correlation to the reference and examined the variability across TRF morphologies.

For statistical comparison at the group level, Wilcoxon signed-rank tests were used to assess differences in all three metrics between CORSICA at its optimal inclusion rate of 98.28% and no artifact rejection. Rank-biserial correlation (RBC) is reported as effect-size and all *p*-values were FDR-corrected for multiple comparisons (Benjamini-Hochberg procedure, *n* = 3).

### 2.8 Software

To make the developed method easily adoptable by other researchers, we developed a Python package that implements the method. The Python package can be downloaded from https://pypi.org/project/corsica-ci/.

## 3 Results

### 3.1 Artifact characterization

To characterize how artifacts produced by the CI manifest in EEG recordings and how they compare to the neural components, we analyzed individual ICs of the EEG data. Fig. 2 shows three exemplary components, characterized by their topography, cross-correlation with the audio waveform, their TRFs for the speech envelope, and their power spectrum. Next to one typical artifactual component and one typical brain component, we also chose a representative mixture component to demonstrate that some components combine features from both ends of the spectrum.

**Figure 2:**
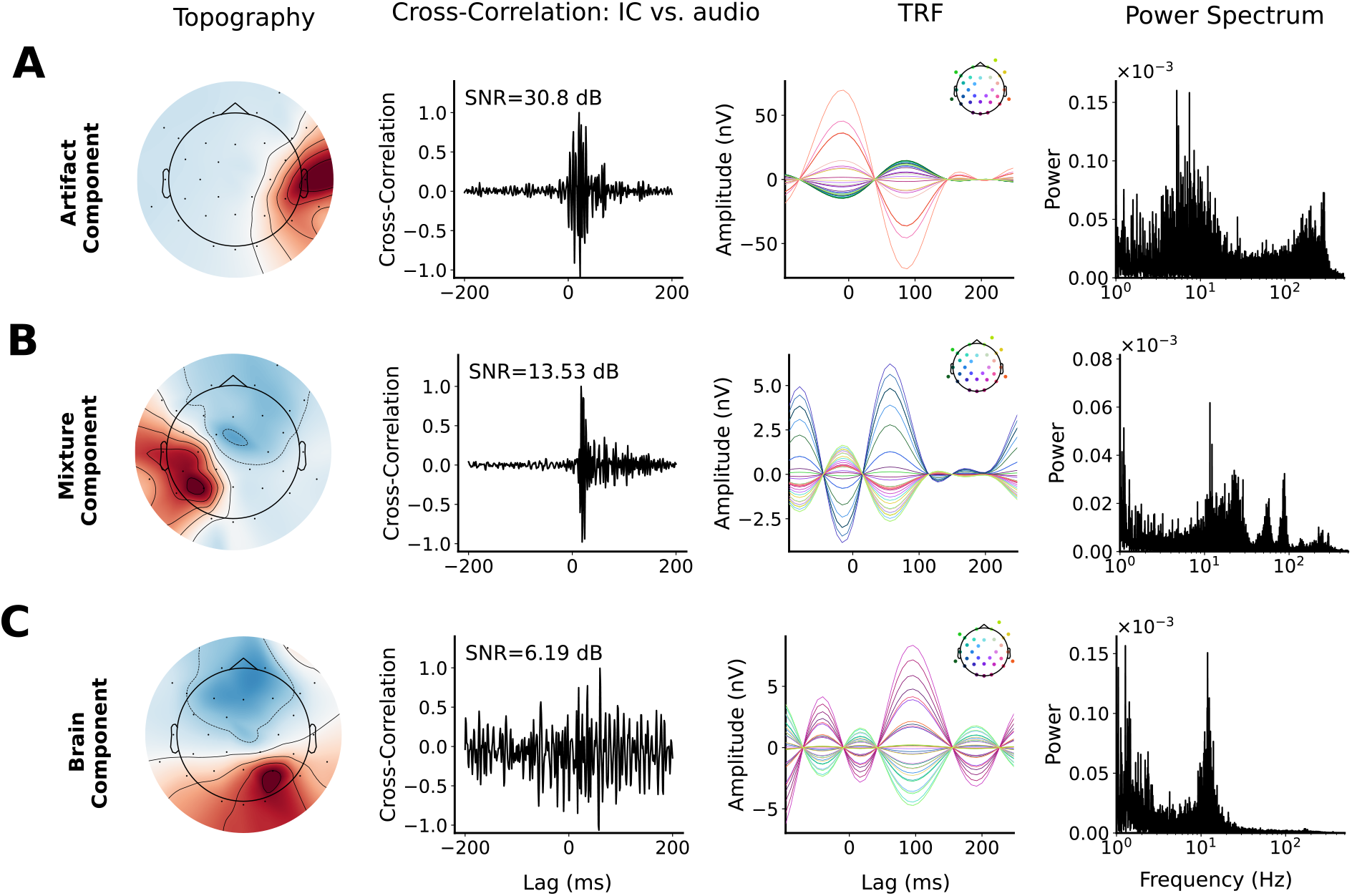
Properties of three exemplary ICs. **A**, An IC with an exemplary CI artifact exhibits an asymmetric topography centered near the right ear, lacking a distinct dipole structure. The cross-correlation with the audio reveals a peak with a high SNR of 30.8 dB at around 0 ms latency. This is also reflected in a peak in the TRF for the speech envelope: this peak is driven by the same lateralized channels that dominate the topography. The power spectrum is characterized by a bimodal distribution, with dominant power around 10 Hz and a secondary high-frequency cluster between 150–300 Hz. **B**, The mixture component shows its strongest pole near the left ear and a second pole in the fronto-central region. While the cross-correlation function reveals a peak at a delay of around 0 ms, the SNR within the proposed search window is significantly lower (13.53 dB) than in purely artifactual components. Similarly, the TRF peak at 0 ms is reduced by an order of magnitude to *−*3 nV. The power spectrum maintains a peak around 100 Hz, but high-frequency power is markedly diminished compared to the artifact component. **C**, The brain component exhibits a clear dipolar topography and lacks a distinct correlation peak with the audio signal. Its TRF is characterized by a physiologically plausible peak around 100 ms. The power spectrum is bimodal, showing expected peaks in the delta (1–4 Hz) and alpha (8–13 Hz) bands, followed by a sharp high-frequency cutof

#### CI artifact component

A typical CI artifact, as depicted in Fig. 2A, is marked by a strongly lateralized topography centered around the position of either of the implants, as these are the positions where the artifacts originate. The topography does not show a dipolar structure. The cross-correlation function of the IC’s timeseries with the audio waveform depicts a sharp peak, with a very high SNR of 30.8 dB in the proposed search window between −5 ms and 12 ms around the onset. This peak reflects the high degree of correspondence between the CI artifact and the audio stimulus. Similar to the topography, the TRF is dominated by a few channels near the right ear that exhibit large amplitude magnitudes at 0 ms. The distribution of power across frequencies is bimodal. Beyond a first peak around 10 Hz, high-frequency components between 150 and 350 Hz contribute substantially to the signal’s power, which are consistent with common stimulation strategies, where frequencies up to 300 Hz are used for apical stimulation [3, 33].

#### Mixture component

As the blind-source-separation rarely achieved a clear-cut separation of artifacts and neural components, we observed several mixture components, which we understand as linear combinations of neural activity and artifactual data. Fig. 2B illustrates a representative example: while the topography is dominated by lateralized activity near the left implant, a secondary fronto-central pole emerges, suggesting a neural contribution. The cross-correlation function maintains a distinct peak. However, the SNR within the search window results in a lower value of 13.5 dB. While the TRF also depicts a substantial response near 0 ms, the amplitude is significantly reduced compared to the artifact component to *−*3 nV. Furthermore, channels from the entire montage contribute to the TRF. Finally, the spectral profile displays a pronounced decay between 1 and 10 Hz with a clear 12 Hz alpha peak, yet it fails to taper at higher frequencies. Instead, we observe persistent power around 100 Hz, likely caused by the CI stimulation.

#### Brain component

Fig. 2C depicts a robustly isolated neural component. Its topography is characterized by a clear dipolar distribution, with poles situated in the fronto-central and slightly right-lateralized posterior regions. The analysis of the temporal correlation with the presented audio confirms the neural origin, as no distinct peak emerges at 0 ms. Similarly, the TRF has no significant peak near 0 ms latency. Instead, the TRF peaks at a physiologically plausible latency of 100 ms with a 7 nV magnitude, and a spatially broad distribution across channels. Lastly, the power spectrum displays a canonical profile, featuring prominent energy in the delta (1–4 Hz) and alpha (8–13 Hz) bands, and a sharp drop of power above 20 Hz.

### 3.2 Qualitative evaluation of artifact detection methods

The TRFs for the preprocessed EEG data – with ICs of physiological artifacts already removed – still showed large CI artifacts, both for the attended and the distractor speaker (Fig. 3A). These artifacts manifested as large magnitudes of a few EEG channels with peaks around the delay of 0 ms. Especially channels located near the CIs, such as channels TP9 and T7 in our IC example in Fig. 2A, exhibited the elevated magnitudes. This demonstrated that CI artifacts survived the removal of physiological artifacts with ICLabel.

**Figure 3:**
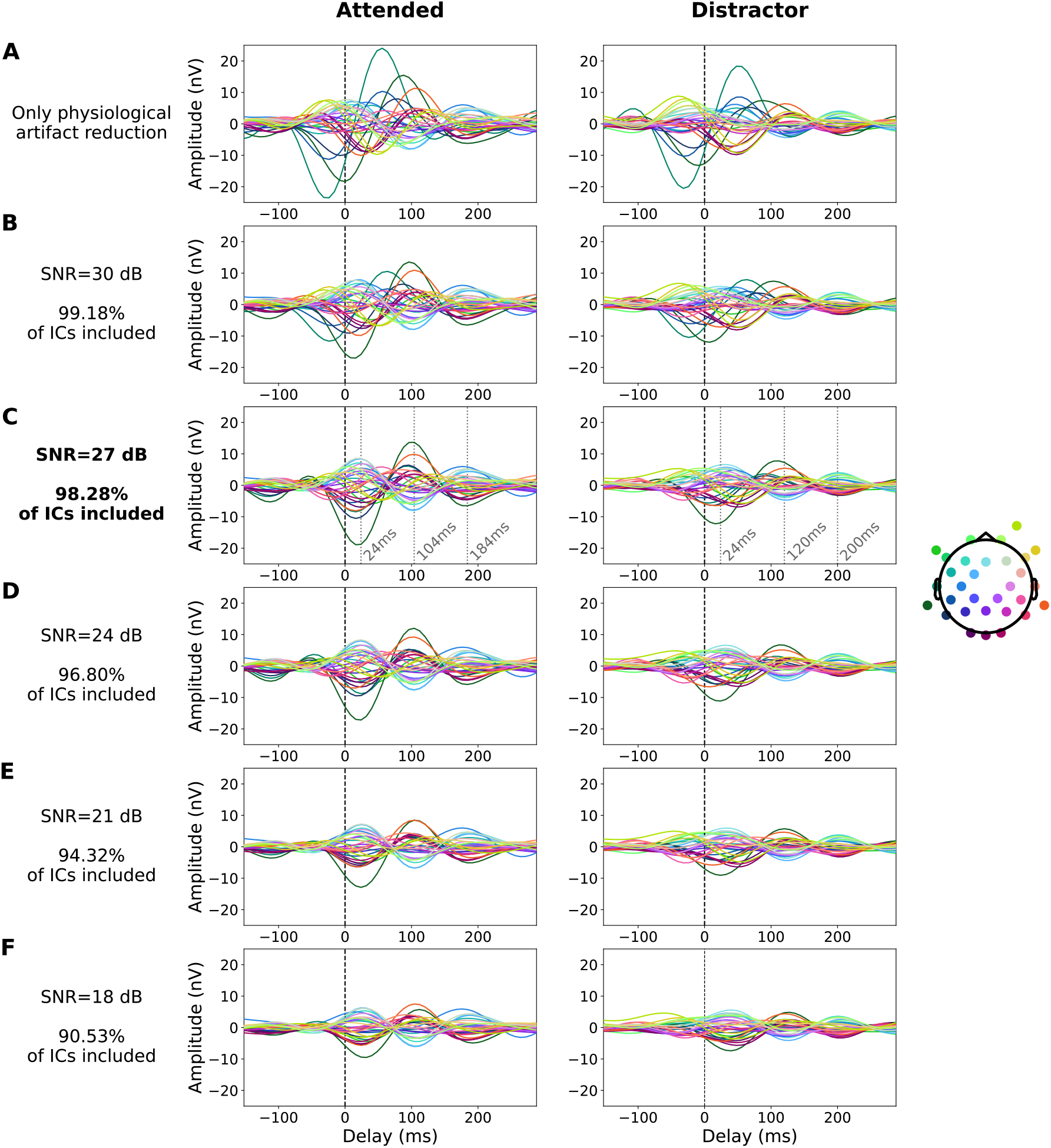
Grand average TRFs resulting from CORSICA. TRFs were computed for different SNR thresholds for attended and distractor stimuli. **A**, The baseline TRF, where only physiological artifacts were removed, still exhibits significant CI artifacts. **B - F**, The highest SNR threshold at which CI artifacts are no longer visible for the attended and distractor speaker is 27 dB SNR (**C**). For the attended talker, the TRFs then peak at 24 ms, 104 ms, and 184 ms, while the peak delays are 24 ms, 120 ms, and 200 ms for the distractor.

We therefore applied CORSICA and the TRF-based benchmark method (TRF_ICA_) to the preprocessed EEG data. We varied the tunable SNR threshold such that the data inclusion rates ranged from 50% to 100%, in order to examine the influence of this parameter on the resulting TRFs. For CORSICA, the SNR threshold was varied from 6 dB to 32 dB in 1 dB increments. At each step, the percentage of included data was calculated as the proportion of retained ICs relative to the total IC count. The SNR thresholds for TRF_ICA_ were chosen to yield comparable data inclusion rates, enabling a fair comparison between the two methods. The TRF_ICA_ threshold was accordingly varied from 5 dB to 25 dB in 1 dB increments.

For both methods, TRFs were computed separately at each SNR threshold for the attended and distractor speakers. Panels B–F in Fig. 3 present TRFs at selected SNR thresholds for CORSICA, while Fig. 4 (Panels B–F) shows the corresponding TRFs for TRF_ICA_ at comparable data inclusion levels.

**Figure 4:**
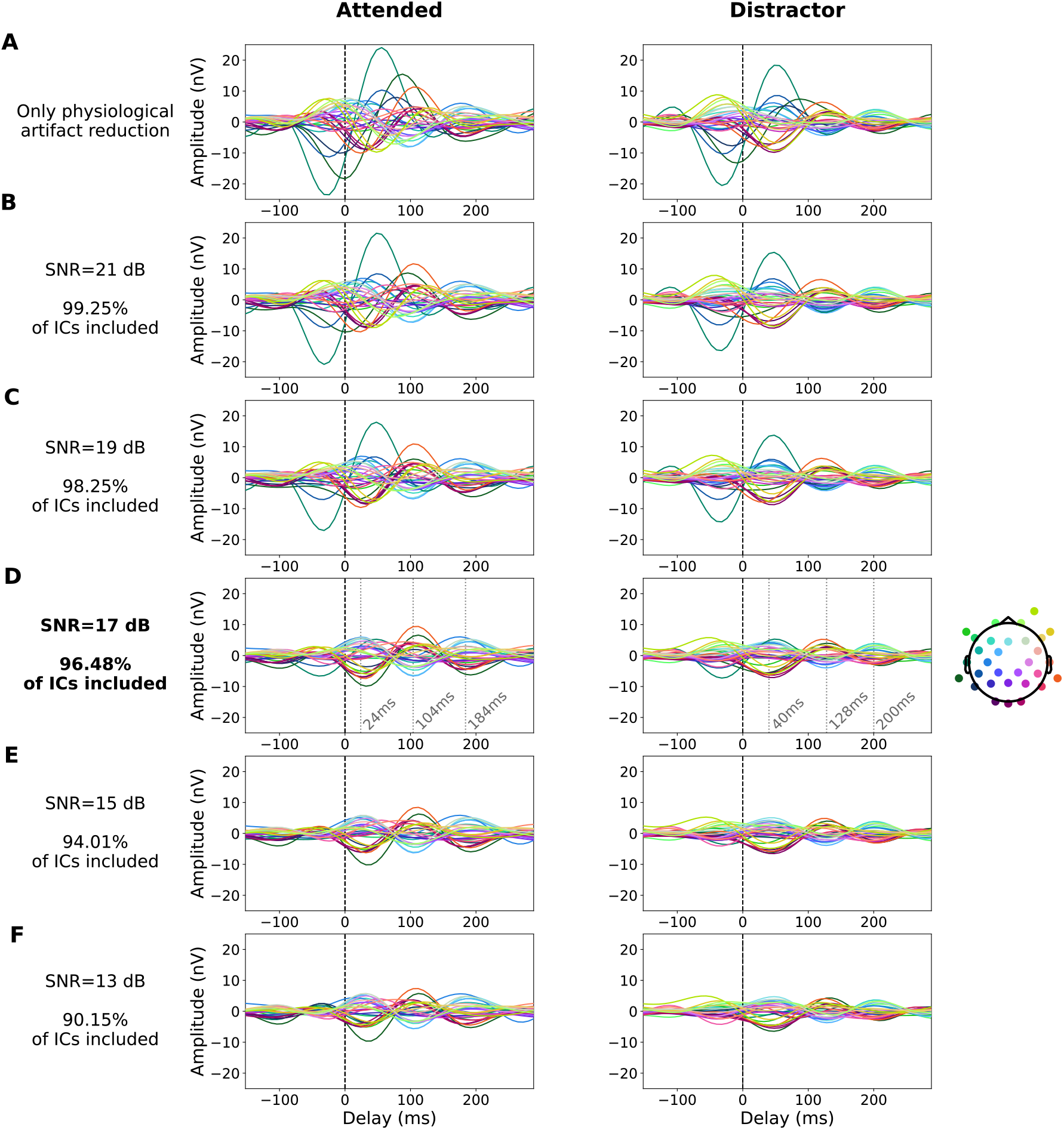
Grand average TRFs from the TRF_ICA_ approach for CI artifact removal. TRFs were computed for different SNR thresholds for attended and distractor stimuli. **A**, Baseline TRF with physiological artifact removal only. **B - F**, TRFs for decreasing SNR thresholds. All but one visible CI artifacts disappeared at a SNR threshold of 17 dB (**D**) and lower. The SNR threshold of 17 dB was thus determined as the optimal one. Grey dotted lines indicate the peak latencies at 24 ms, 104 ms, and 184 ms for the attended speaker. The distractor TRF peaks emerge at latencies of 40 ms, 128 ms, and 200 ms.

#### CORSICA

When applying CORSICA, the artifact contamination of the TRFs was already strongly reduced when only 0.82% of the data were excluded (SNR threshold of 30 dB), with a clear reduction in the magnitude of the two prominent channels (Fig. 3B-F). As more data was excluded (lower SNR thresholds), the peaks of typical artifact-contaminated channels such as TP9 and T7 shifted toward longer and thus more expected latencies, suggesting effective CI artifact suppression. At the SNR threshold of 27 dB, CI artifacts were greatly attenuated, while a large percentage of the EEG data (98.28% of all ICs) was preserved. The prominent peaks then emerged at physiologically plausible latencies of 24 ms, 104 ms, and 184 ms. Therefore, we selected 27 dB as the optimal SNR threshold for CORSICA. For lower SNR thresholds, the attended TRFs remained qualitatively similar with decreasing amplitudes.

The distractor TRFs yielded similar results, as decreasing SNR thresholds led to reduced magnitudes and longer peak latencies. Overall, the TRF magnitudes were generally smaller in the distractor condition compared to the attended condition. The distractor TRF peaked at later latencies (24 ms, 120 ms, and 200 ms) compared to the attended condition as expected [9].

#### TRF-based artifact detection

Figure 4 shows the TRFs at different SNR thresholds for the TRF-based artifact reduction method. Inspecting the TRFs at the 19 dB threshold (Figure 4C), where the same amount of data as the optimal threshold for the correlation-based approach is preserved, we still observe a prominent peak in both the attended and distractor TRF before stimulus onset. The early latency and the location of the peaking channel T7 near the left ear are both indicative of a persistent artifact. CI artifacts decreased notably and were barely visible at an SNR threshold of 17 dB (Figure 4D) or lower. We thus evaluated the optimal threshold as 17 dB, where 96.48% of data was preserved. The distractor peak latencies (40 ms, 128 ms, and 200 ms) were slightly delayed compared to the attended peaks (24 ms, 104 ms, 184 ms).

#### Peak amplitudes

Since more data was excluded in the TRF_ICA_ approach to reach a satisfactory TRF than for CORSICA, the TRF amplitudes appeared lower in the former than in the latter case. To quantify this amplitude reduction, we calculated the absolute value of each channel’s TRF coefficient and then took the average across all channels. We found average peak amplitudes of 4.9 nV, 3.9 nV, and 2.5 nV for the attended TRF for the CORSICA method at 27 dB SNR threshold, and 3.3 nV, 2.8 nV, and 2.4 nV for the TRF_ICA_ approach at 17 dB SNR threshold. The numerical analysis thus supported the visual impression.

#### Comparison to SOBI blind source separation

We next investigated the impact of the source separation method on CI artifact removal, comparing ICA against second-order blind source separation (SOBI) as the backbone for artifact rejection. To this end, we repeated the correlation-based analysis using SOBI instead of InfoMax ICA to identify independent EEG components. Those components were subsequently evaluated for CI artifacts using the correlation based method to quantify the SNR. Given that CORSICA outperformed TRF_ICA_ on ICA-derived components, TRF_SOBI_ was not considered for the SOBI comparison.

The resulting TRFs from CORSICA and the correlation-based method using SOBI source-separation (corr_SOBI_ are compared in Fig. 5. The optimal SNR threshold for CORSICA of 27 dB, corresponding to a data retention of 98.12%, served as the baseline. For corr_SOBI_, an equivalent IC retention rate required a notably higher SNR threshold of 33.0 dB, suggesting that SOBI yielded less separable artifact components than ICA.

**Figure 5:**
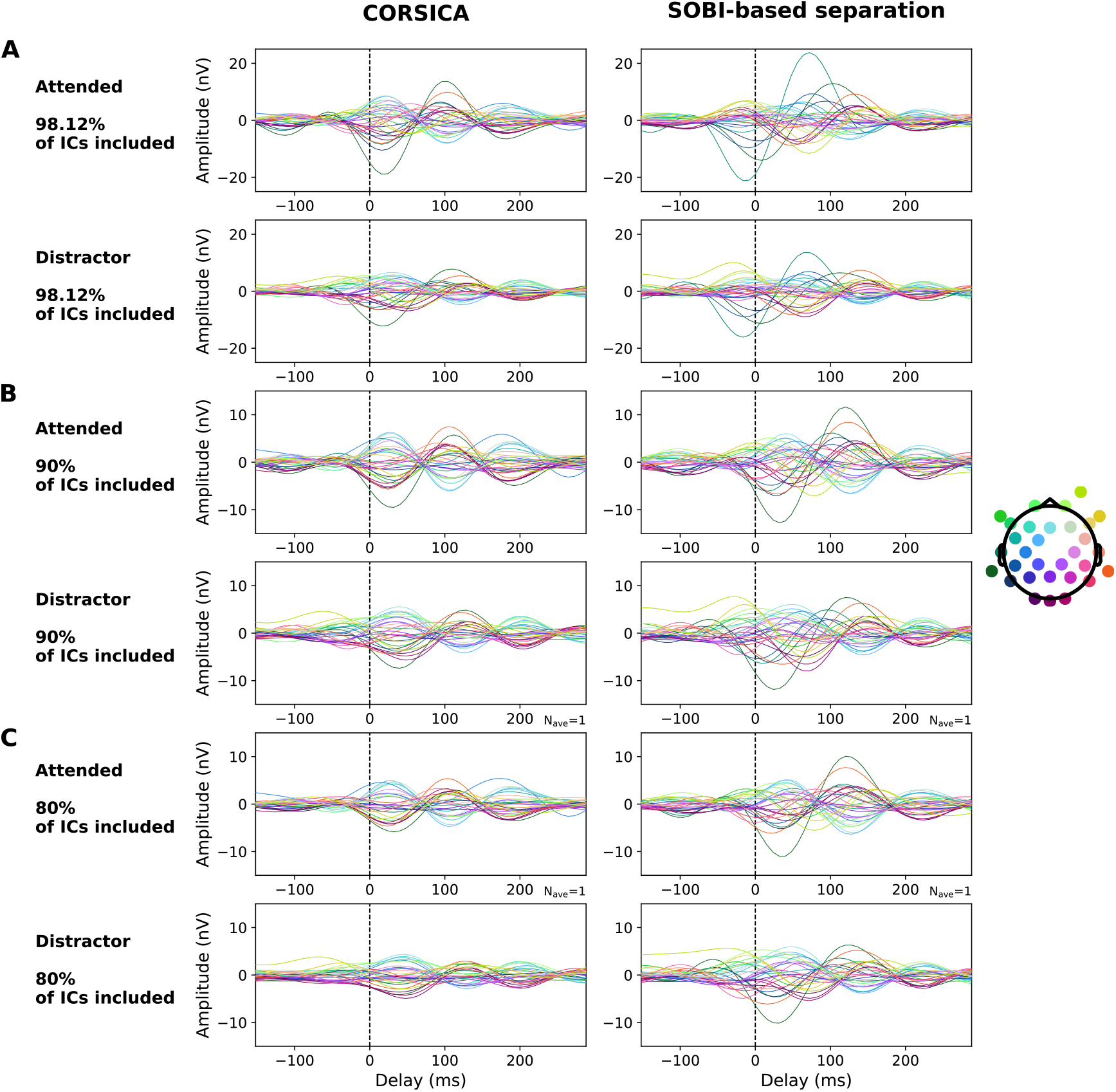
Grand average TRFs using the same correlation-based artifact rejection approach but involving two different source-separation methods. Left column: corr_ICA_, which is used for CORSICA method. Here, sources were separated into independent components using InfoMax ICA. Right column: corr_SOBI_ method where sources are separated by second-order blind identification (SOBI). The percentage of included ICs is identical within each row to facilitate comparability across methods. The method based on SOBI requires a larger portion of excluded ICs to achieve successful removal of the CI-artifacts.

While the artifacts were largely attenuated for the ICA-based separation, prominent CI-artifacts remained visible for the SOBI-based separation, for both the attended and distractor TRF (Fig. 5 A). The physiological plausibility of the TRF based on corr_SOBI_ was substantially improved when including only 90% of ICs (SNR threshold=27.3 dB, Fig. 5 B). Peaks in the negative domain had then vanished, and the artifact’s amplitude had decreased. However, a single dominant channel from the left hemisphere remained visible for both the attended and ignored TRF obtained from corr_SOBI_.

In contrast, the TRF obtained from CORSICA appeared entirely artifact-free at this stage. Note that for Fig. 5C-F, the scale has been reduced to −15-15 nV to enable a clear visibility of the TRFs’ shapes. A satisfactory artifact attenuation using the corr_SOBI_ method was achieved at 80% of ICs included (SNR threshold = 22.9 dB, Fig. 5 C). Both the attended and distractor TRF were free from obvious artifacts, as the impact of single channels had further decreased. However, as more components had to be excluded to ensure acceptable signal quality, the signal power was reduced, which translated to lower TRF amplitudes than for CORSICA at 98.12% IC inclusion.

### 3.3 Metric-based comparison of artifact reduction methods

Figure 6 compares the TRFs obtained from the three employed artifact rejection methods, CORSICA, corr_SOBI_ and TRF_ICA_, across varying levels of data inclusion based on Pearson correlation, the averaged latency error of TRF peaks compared to the reference, and the averaged amplitude error of the TRF peaks. Given that a ground truth TRF is lacking, we used the TRF obtained from the attended speaker with the CORSICA method at the optimal SNR threshold of 27 dB (98.28% data retained) as a reference. This TRF was previously identified as optimal through visual inspection. Note that we applied a nonlinear power-law scaling (exponent=2.5) to the x-axis to improve the visibility of changes at high levels of data inclusion. SNR thresholds were varied in integer steps, which correspond to unequal increases in data retention, resulting in unevenly spaced data points along the *x*-axis.

**Figure 6:**
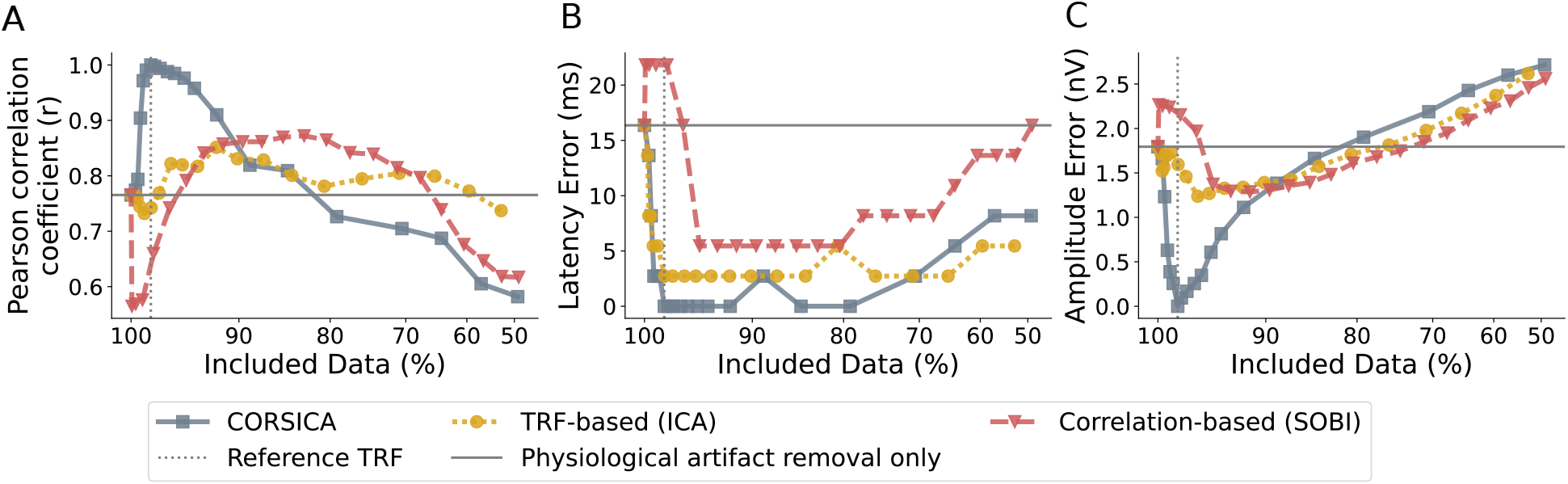
Metric-based comparison across CORSICA and the TRF_ICA_, and corr_SOBI_ benchmarks for CI artifact removal. The TRF obtained from CORSICA at an optimal SNR threshold of 27 dB was employed as a reference. The corresponding percentage of data inclusion of 98.12% is marked as a vertical dashed grey line. The x-axis has been nonlin-early transformed using a power-law scaling (exponent of 2.5) to enhance the visibility of variations at high levels of data inclusion. The horizontal grey line indicates the baseline of each metric for data where only physiological artifact removal was applied. **A**, Pearson correlation between the different TRFs and the reference TRF, averaged across EEG channels (higher is better). Both ICA-based methods first show an increase in correlation with the reference TRF, and reach a maximum around the reference ratio of data inclusion. The SOBI-based approach first exhibits a decrease in correlation to the reference TRF and reaches its maximum when 90% of independent components are included. **B**, Latency error (lower is better). Both ICA-based methods show a sharp improvement when 2% of ICs are rejected. A minimum latency error of 3 ms is reached by the TRF-based approach. The SOBI-based correlation method reaches the lowest latency difference of 5 ms to the reference TRF between 94% and 80% data inclusion. **C**, Amplitude error (lower is better). The TRF-based method reached its minimum slightly after the reference TRF at 96% data inclusion. The SOBI-based method reached its minimum only when 82% of independent components are included. All methods exhibit a substantial increase in amplitude error, when more than 20% of components are rejected.

#### Pearson correlation

At very high SNR thresholds, where almost 100% of the data are retained, both ICA-based methods for CI artifact removal exhibited only medium values of channel-wise Pearson correlation with the reference TRF (Fig. 6A). This reflected the presence of residual CI-induced artifacts, as also observed in the corresponding TRFs (see Fig. 3B and Fig. 4B,C). As the rejection threshold decreased, such that more components were rejected, the correlation first improved. CORSICA yielded higher correlation values than the TRF_ICA_ approach, which was expected, as the perfect correlation value of *r* = 1 was necessarily reached when the reference TRF was correlated with itself. The TRF_ICA_ method achieved its maximum correlation with the reference TRF (*r* = 0.85) also at 96% data retention, which is consistent with the visual inspection. The overall strong agreement between the two methods (maximal correlation value of *r* = 0.85) supported that both approaches largely eliminated the same ICs, and demonstrated that key TRF features were reproducibly captured with both artifact removal methods. In contrast, with decreasing SNR threshold, the corr_SOBI_ approach first suffered from a decrease in correlation to the reference TRF as obtained by CORSICA, possibly due to the different source-separation approach. However, stronger artifact rejection then lead to a substantial increase in correlation, and a maximum of *r* = 0.86 was reached when 90% of ICs were included. Hence, the corr_SOBI_ reached a slightly higher maximal Pearson correlation with the reference TRF, obtained by applying CORSICA, compared to the TRF_ICA_ approach.

#### Latency error

Fig. 6B reports the latency error and shows that the mean absolute latency error for the CORSICA approach reached zero at 98.12% data inclusion, as expected for the reference threshold. Noteworthy, however, is that the latency error remained zero even when removing about 20% of the data. The TRF_ICA_ method followed a similar trend, reaching 3 ms latency error at 98.12% data inclusion and staying there until up to 35% of the data were excluded. In both methods, the latency error increased when more than 20% of the data were excluded, consistent with an over-rejection of components and the following signal quality loss. After an initial increase in latency error to 22 ms, the corr_SOBI_ method dropped to a 5 ms latency error between 94 and 80% of included ICs.

#### Amplitude error

As reported in Fig. 6C, the amplitude error of CORSICA reached its inherent minimum at 98.28% data inclusion, with sharp increases observed on either side of this optimum. The TRF-based method mirrored this trajectory, similarly peaking in performance at 96.48% inclusion with a minimum amplitude error of 1.3 nV. Similar to the other metrics, the SOBI-based method first exhibited a slight deterioration before the amplitude error fell off. The minimum latency error of 1.3 nV was reached at 90% data inclusion, followed by a steady increase with stronger artifact rejection.

Across the three metrics, we observed that the TRF_ICA_ method reached its optimum consistent with visual inspection around 96% data inclusion, further supporting this threshold as optimal. Consistent with the visual inspection, the corr_SOBI_ approach reached its optimum with respect to the reported metrics later than the other methods. This suggests that, when relying on temporal correlation to the audio signal as an exclusion criterion, more independent components must be excluded when using SOBI compared to Infomax ICA.

### 3.4 Subject-level analysis

The preceeding analysis targeted the population behaviour, with results computed from the EEG data of all CI patients combined. To assess how well CORSICA worked on the level of individual subjects, and how variable the results at this level were, we also computed performance metrics and TRFs for single subjects (7).

In Fig. 7A, we report Pearson’s correlation coefficient, latency and amplitude error of the TRFs on the level of individual subjects, as compared to the reference TRF. Additionally, we report the corresponding metrics for three individual subjects, one with good CORSICA performance, another with medium performance, and a third with poor performance.

**Figure 7:**
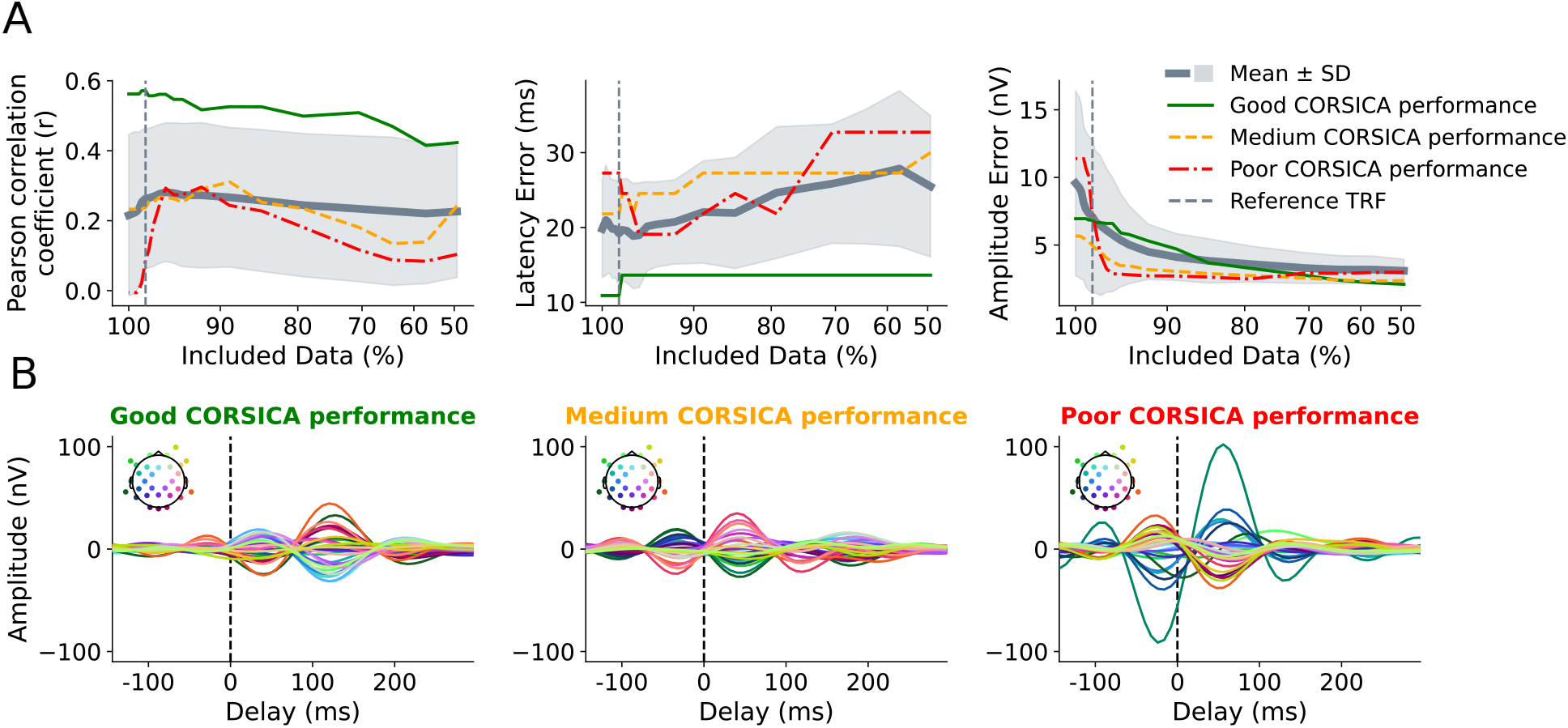
Subject-level analysis of TRFs obtained through CORSICA. **A**, Evaluation of subject-level TRFs through different metrics (Pearson correlation coefficient, latency error, and amplitude error), relative to the grand-average reference TRF. The bold grey line indicates the mean across subjects, and the grey-shaded area denotes one standard-deviation. Results from three subjects with good, medium and poor CORSICA performance are shown as green, yellow and red lines, respectively. The vertical dashed line indicates the optimal value of 98.28% data inclusion determined from the grand average. The mean of Pearson correlation coefficient peaks at 96% data inclusion before declining gradually at lower inclusion rates. The latency error reaches a minimum at 96% data inclusion, which barely differs from the baseline, before rising above the baseline value, while the amplitude error decreases monotonically with decreasing data inclusion. **B**, TRFs of the three exemplary subjects from panels A, at 98.25% data inclusion. The TRFs on the left show a clean dominant peak around 120 ms with no visible artifact residuals. The TRFs of the middle panel exhibit a peak at negative latencies, likely reflecting residual CI contamination. The TRFs of the right panel remain dominated by a few channels near the left implant, indicating insufficient artifact suppression.

#### Pearson correlation coefficient

At the level of individual subjects, Pearson correlation coefficients were overall lower than in the grand-average analysis, reflecting the inherent noisiness of individual TRFs. A clear increase with artifact rejection was nonetheless observable, with Pearson correlation coefficients peaking at an optimal inclusion rate of 96.16%, which was slightly lower than the grand-average optimum of 98.28%. This discrepancy reflects a difference between the mean of the individual correlation coefficients and the correlation coefficient of the grand average. Such a difference can arise from the nonlinearities involved in the ridge regression as well in the computation of Pearson correlation coefficients.

At high data exclusion rates, Pearson correlation coefficients declined and converged toward the no-rejection baseline. Comparing no artifact rejection to CORSICA at 98.28% data inclusion, a Wilcoxon signed-rank test revealed a significant improvement in the correlation coefficients, with a large group-level effect (*W* = 18.0, *RBC* = 0.789, *p* = 0.0053).

#### Latency error

Without artifact rejection, the mean latency error on the level of individual subjects was comparable to the grand average (20 ms). However, unlike the grand-average, no systematic reduction in latency error was observed across SNR thresholds, indicating that TRF peak latencies only become reliable after averaging across subjects. Accordingly, the Wilcoxon signed-rank test revealed no significant improvement at 98.28% data inclusion relative to no artifact rejection (*W* = 6.0, *RBC* = *−*0.2, *p* = 0.787). Consistent with Pearson correlation coefficient, the subject with good CORSICA performance achieved the lowest latency error, while the subjects with medium and poor CORSICA performance exhibited substantially larger errors of 21 and 29 ms, respectively.

#### Amplitude error

The baseline amplitude error (no artifact rejection) at the level of individual subjects was substantially larger than in the grand-average analysis (10 nV vs. 2 nV). This presumably reflects the observation that TRFs at the level of individual subjects can exhibit peaks at different latencies from those of the grand average TRF, which contribute to the amplitude error, while many of these peaks are suppressed when averaging across subjects.

The amplitude error showed a clearest trend with increasing artifact rejection, decreasing monotonically towards a value of about 2.9 nV. Accordingly, the Wilcoxon signed-rank test revealed a highly significant reduction in amplitude error at 98.28% data inclusion, as compared to no artifact rejection, with a large effect size (*W* = 2.0, *RBC* =-0.97, *p <* 0.001). Interestingly, the subject with the good CORSICA performance exhibited a larger amplitude error than the subject with medium performance, although better (at 98.28% data inclusion) than the subject with medium performance.

#### Individual TRFs

Fig. 7B shows individual TRFs obtained with CORSICA at 98.28% data inclusion (vertical dashed line in panels A). The TRFs of the subject with good CORSICA performance show no artifactual response near 0 ms latency, but instead a first peak at about 40 ms delay and a second larger peak around 120 ms that are clearly of neural origin. The subject with medium CORSICA performance exhibits TRFs with a residual response at negative delays, indicative of incomplete artifact suppression, alongside neural responses at around 40 ms and 200 ms. The TRFs of the subject with poor CORSICA performance remain dominated by the CI artifact, driven by several left-lateralized channels, with the magnitude of one channel clearly exceeding the neural responses seen in the other examples.

## 4 Discussion

In this study, we introduced CORSICA, a reproducible and fully objective method for CI artifact rejection in EEG responses to continuous speech. We benchmarked the method thoroughly against two alternatives: a TRF-based rejection criterion and SOBI as an alternative source separation backbone. By systematically evaluating the resulting TRFs across a range of SNR thresholds, we demonstrated that CORSICA effectively suppresses CI artifacts while retaining more than 98% of independent components, leaving neural activity largely intact. These results establish the temporal correspondence between the stimulus and the CI artifact as a robust and sufficient criterion for artifact identification, and validate CORSICA as a practical pre-processing standard for speech-evoked EEG in CI users.

This work closes a methodological gap in CI research, as previous approaches are either ineffective for continuous stimuli [14] or rely heavily on manual component labeling, which is time-consuming and lacks reproducibility [7, 17]. As interest in EEG recordings from CI users continues to grow, CORSICA offers the research community an objective, validated, and parameter-transparent solution for CI artifact reduction.

### 4.1 Method comparison

CORSICA identifies CI artifacts through the cross-correlation between the audio stimulus and each IC time series, penalizing components that yield peaks near 0 ms delay — a signature of the tight temporal coupling between the stimulus and the resulting artifact. The TRF-based benchmark (TRF_ICA_) operates on the same principle but uses the TRF of each IC with respect to the speech envelope instead of the broadband cross-correlation.

Although both approaches effectively suppressed CI artifacts and recovered physiologically plausible TRFs, CORSICA proved superior. Rejecting only 1.78% of ICs was sufficient for robust artifact suppression, compared to 3.52% for TRF_ICA_, and the TRFs resulting from CORSICA exhibited larger peak amplitudes. The advantage of CORSICA may stem from its use of broadband signal content up to 300 Hz, whereas TRF_ICA_ relies on the speech envelope in the much narrower 1–8 Hz range, potentially providing a less discriminative artifact signature.

Replacing ICA with SOBI as the source separation backbone (corr_SOBI_) also achieved robust artifact attenuation, but required the removal of a substantially larger proportion of ICs. This was consistent across both visual inspection, where 20% data exclusion was deemed appropriate, and the metric-based analysis, where the optimal operating point for corr_SOBI_ fell between 10% and 20% exclusion — well above the 1.78% sufficient for CORSICA.

### 4.2 Participant variability

The subject-level analysis revealed substantial inter-individual variability in TRFs, similar to what is known from typical hearing individuals [36]. However, different neural processing and CI artifacts amplified this effect even further.

Statistically, CORSICA significantly improved both Pearson correlation and amplitude error relative to no artifact rejection, with a particularly large effect for amplitude error (RBC = *−*0.97), confirming its effectiveness in suppressing high-amplitude CI artifacts. The complementary behavior of the metrics underlines the importance of evaluating multiple quality measures: amplitude error decreased monotonically with more aggressive rejection, whereas Pearson correlation peaked at 96.16% and declined beyond this point, demonstrating that overly aggressive rejection degrades TRF morphology.

Latency error was not significantly improved at the subject level, suggesting that it is particularly susceptible to individual TRF noise and only becomes reliable after across-subject averaging. Finally, the variability across the three exemplary subjects shown in Fig. 7 suggests that a subject-specific SNR threshold could improve outcomes for participants with pronounced residual artifacts, at the cost of reduced pre-processing comparability across subjects. Such subject-specific preprocessing constitutes an interesting direction for future work.

### 4.3 Temporal characteristics

Using temporal characteristics for artifact detection has previously been applied by Paul et al., who identified artifact components by visually comparing IC activations with the audio stimulus waveform [7, 8]. When the component’s time series mirrored the temporal dynamics of the stimulus, those components were classified as artifacts. Our work further supports the important role of temporal alignment to identify CI artifacts. The methods employed in the current study take previous approaches one step further by introducing a quantitative SNR value to measure the strength of temporal alignment between the stimulus and the independent component. We note that the semi-automated method by Viola et al. [14] identifies CI artifacts by their sharp onset and offset with stimulus transition for syllable stimuli. The methods that we propose hear may be viewed as a generalization of this approach for EEG responses to continuous audio stimuli, where a clear temporal separation between stimulus onset and offset is lacking.

### 4.4 Spatial characteristics

CI artifacts have previously routinely been identified through manual inspection of the spatial topographies of independent components [7, 13, 37]. CI-related components typically exhibit strong activity localized near one of the implant sides, as depicted in Fig. 2 A. However, the transition to components that contain a mixture of neural signal and artifact, as shown in Fig. 2 B, can be hard to detect. To date, no objective method to classify CI-artifacts to continuous stimuli based on their topography is available. Moreover, the topography alone cannot indicate the strength of an artifactual component. Therefore, when relying on manual labeling of components, temporal and spectral characteristics should also be considered.

### 4.5 Spectral characteristics

The power spectrum of the exemplary CI-artifact in Fig. 2 A showed significant contributions at frequencies between 150 Hz and 300 Hz, which is the frequency range typically used for apical stimulation [33]. Those high-frequency components are consistent with prior results on CI artifacts in electrically-evoked auditory steady-state response measurements [38]. In contrast, neural components exhibit a 1/f power decay and thus have only negligible power above 100 Hz (Fig. 2 C). Previously, only Kim et al. [17] considered spectral information to identify CI artifacts to speech stimuli by visually identifying spectral anomalies in the power spectrum of the EEG up to 50 Hz. As a result of their low cut-off frequency in the preprocessing, they likely have missed some high-frequency components of the CI artifact. The method was only validated in four individuals and relied on manual inspection. Nevertheless, the success of our correlation-based method, incorporating high-frequency data, and the high-frequency content in the exemplary artifact suggest that spectral characteristics are a useful marker for identifying CI-artifacts.

### 4.6 Limitations

One limitation of our study is the reliance on a manually defined ground truth for evaluating the TRF metrics. The reference TRF was chosen based on visual inspection of the TRFs and served as the basis for the metric-based evaluation of the two methods. While this approach was necessary, as no artifact-free dataset was available, it introduced a degree of subjectivity and bias in the evaluation. As such, future work may benefit from independent validation using externally verified ground truth data. Furthermore, our approaches have not yet been validated on other datasets, as publicly available EEG responses to continuous speech in CI users are currently lacking. Future work is needed to assess the generalizability of the proposed methods to other datasets and to determine how the optimal SNR thresholds may vary under different recording conditions.

We also note that both methods developed here rely on the temporal alignment between the stimulus and the independent component. It should, however, be noted that certain device-related factors, such as blocking capacitors introducing delayed signal variations or RF transmission can lead to signals that do not strictly correlate with the acoustic input. Such effect could give rise to artifacts that are not fully captured by our approach.

### 4.7 Conclusion

We presented CORSICA, a fully objective method for CI artifact reduction in EEG responses to continuous speech. We thoroughly benchmarked this method against a TRF-based alternative rejection criterion and SOBI as an alternative source separation backbone. The developed methods rely on the temporal alignment of CI artifacts with aspects of the speech signal at short temporal delays, exploiting the contrast with neural responses which reflect low-frequency speech features at latencies imposed by the auditory pathway. Our evaluation showed that CORSICA outperformed the TRF-based benchmark, and that using SOBI instead of InfoMax ICA required more ICs to be excluded. Excluding only 1.78% of ICs was sufficient for effective CI artifact suppression, yielding clean TRFs. Our findings support the applicability of CORSICA for artifact reduction in real-world, continuous-stimulus EEG recordings without requiring manual artifact labeling.

## Declaration of generative AI and AI-assisted technologies in the writing process

During the preparation of this work, the authors used ChatGPT and Google Gemini in order to improve readability and language. After using these tools, the authors reviewed and edited the content as needed and take full responsibility for the content of the published article.

## Data availability

The data used for this study is publicly available under https://doi.org/10.5281/zenodo.17952844.

